# Taxon-specific shifts in bacterial and archaeal transcription of dissolved organic matter cycling genes in a stratified fjord

**DOI:** 10.1101/2021.06.02.446852

**Authors:** Benjamin Pontiller, Clara Pérez-Martínez, Carina Bunse, Christofer M.G. Osbeck, José M. González, Daniel Lundin, Jarone Pinhassi

## Abstract

A considerable fraction of organic matter derived from photosynthesis in the euphotic zone settles into the ocean’s interior, and under way is degraded by diverse microbial consortia that utilize a suite of extracellular enzymes and membrane transporters. Still, the molecular details that regulate carbon cycling across depths remain little explored. As stratification in fjords has made them attractive models to explore patterns in biological oceanography, we here analyzed bacterial and archaeal transcription in samples from five depth layers in the Gullmar Fjord, Sweden. Transcriptional variation over depth correlated with gradients in chlorophyll *a* and nutrient concentrations. Differences in transcription between sampling dates (summer and early autumn), were strongly correlated with ammonium concentrations, which potentially was linked with a stronger influence of (micro-)zooplankton grazing in summer. Transcriptional investment in carbohydrate-active enzymes (CAZymes) decreased with depth and shifted toward peptidases, partly a result of elevated CAZyme transcription by Flavobacteriales, Cellvibrionales and Synechococcales at 2-25 m and a dominance of peptidase transcription by Alteromonadales and Rhodobacterales from 50 m and down. In particular, CAZymes for chitin, laminarin, and glycogen were important. High levels of transcription of ammonium transporters by Thaumarchaeota at depth (up to 18% of total transcription), along with the genes for ammonia oxidation and CO_2_-fixation, indicated that chemolithoautotrophy contributed to the carbon flux in the fjord. The taxon-specific expression of functional genes for processing of the marine DOM pool and nutrients across depths emphasizes the importance of different microbial foraging mechanisms across spatiotemporal scales for shaping biogeochemical cycles.

**IMPORTANCE:** It is generally recognized that stratification in the ocean strongly influences both the community composition and the distribution of ecological functions of microbial communities, which in turn are expected to shape the biogeochemical cycling of essential elements over depth. Here we used metatranscriptomics analyses to infer molecular detail on the distribution of gene systems central to the utilization of organic matter in a stratified marine system. We thereby uncovered that pronounced shifts in transcription of genes encoding CAZymes, peptidases, and membrane transporters occurred over depth among key prokaryotic orders. This implies that sequential utilization and transformation of organic matter through the water column is a key feature that ultimately influences the efficiency of the biological carbon pump.

## INTRODUCTION

Major portions of the primary production in the photic zone – up to ~40% of the photosynthetically fixed carbon – is transported vertically in the form of particulate organic matter into the ocean’s interior in a process referred to as the biological carbon pump (1). This sinking organic matter is degraded by heterotrophic bacteria via extracellular enzymes that remineralize large proportions to carbon dioxide (2). Sinking particles cross steep gradients in light, temperature, nutrients, and hydrostatic pressure (3). Commonly, density gradients limit mixing between water masses, which disrupts the connectivity of microbial communities and nutrient fluxes. Stratification thereby strongly influences both the microbial community composition and the ecological function of these communities (4). This is for example visible in the replacement of phototrophy genes dominating in surface waters by chemolithoautotrophy genes at depth (5–9). This suggests that a pronounced variability in the genetic repertoire of bacteria and archaea is involved in the processing and uptake of nutrients and organic matter throughout the water column in the open ocean.

In the ocean, the bulk – community level – extracellular enzymatic hydrolysis rates rapidly decrease from epipelagic to bathypelagic zones, whereas the per-cell rates increase, indicating an increased microbial reliance on high molecular weight dissolved organic matter (HMW-DOM) with depth (10, 11). Actually, the hydrolysis of HMW-DOM is considered the rate-limiting step in the marine carbon cycle (12) and bacteria secrete hydrolytic enzymes to utilize particular organic matter and biopolymers (13), e.g. carbohydrate-active enzymes (CAZymes) and peptidases (PEPs), that cleave HMW-DOM into molecules smaller than ~600 Da that can be transported through the cell membrane (14). A few recent studies applied metagenomics to study the spatial and vertical distribution of CAZyme and PEP genes (15–18). For instance, analysis of 94 metagenome-assembled genomes (MAGs) from the Mediterranean Sea uncovered a pronounced depth-related taxonomic and functional specialization in degradation of polysaccharides dominated by Bacteroidetes, Verrucomicrobia, and Cyanobacteria (18). Zhao and colleagues (2020) applied a multi-omics approach encompassing a broad spatial and vertical coverage, showing that both the abundance and diversity of dissolved enzymes excreted by particle-attached prokaryotes consistently increased from epi- to bathypelagic waters (16). Still, knowledge of the concrete expression of these enzymes by different taxa through the water column is limited.

Fjord ecosystems exhibit pronounced vertical gradients in physicochemical and biological conditions, in qualitative aspects reflecting gradients observed in other stratified coastal and offshore marine waters. Due to the reduced scaling of conditions in space and time, fjords are at times referred to as model oceans (19–21). Kristineberg Marine Research Station, created in 1877 and located in the Gullmar Fjord – a sill fjord on the west coast of Sweden – is one of the oldest marine research stations in the world. The extensive knowledge of the physical, chemical, and biological oceanography in the fjord provides a solid background against which to determine aspects of the microbial oceanography (22). Therefore, to obtain novel mechanistic knowledge of the functional degradation of biopolymers with depth and the microbial taxa that produce the required enzymes, we applied environmental metatranscriptomics to investigate the expression of polymer degrading enzyme systems (i.e., CAZymes and PEPs). As we aimed at studying the combined responses of both degradation and uptake of ecologically important DOM compounds or nutrients in this system, we included membrane transporters in the analysis. We hereby hypothesized that potential divergence in CAZyme, PEP, and transporter expression would be associated with shifts in dominance of transcription levels among taxa across depths.

## MATERIAL & METHODS

### Study site and sampling

This study was conducted in the Gullmar Fjord at station Alsbäck (58°19′22.7″ N; 11°32′49.0″ E) (Fig. 1). The fjord is located on the Swedish west coast approximately 100 km north of Gothenburg. As in other fjords, the Gullmar Fjord has pronounced differences in residence time between the dynamic surface (16 – 40 days) and deep waters (46 days to a year) (23), which ultimately structures the distribution of nutrients and microbiota. We sampled a vertical depth profile spanning from surface water to 100 m depth in July and September 2017. CTD profiles were obtained prior to the selection of the desired depth layers onboard the R/V *Oscar von Sydow*. In July, we sampled water from 5, 15, 50, 75, and 100 m depth, whereas in September we took water from 2, 25, 55, 75, and 100 m depth. The different depth layers were chosen based on CTD casts and represented major transitions in physicochemical parameters (Fig. S1A-D). From all depth layers, we sampled biological duplicates with a 30L Niskin bottle for additional measurements such as nutrients, chlorophyll *a* (Chl *a*), dissolved organic carbon (DOC), and metatranscriptomics. Detailed information is provided in Supplementary File 1.

**Figure 1.**
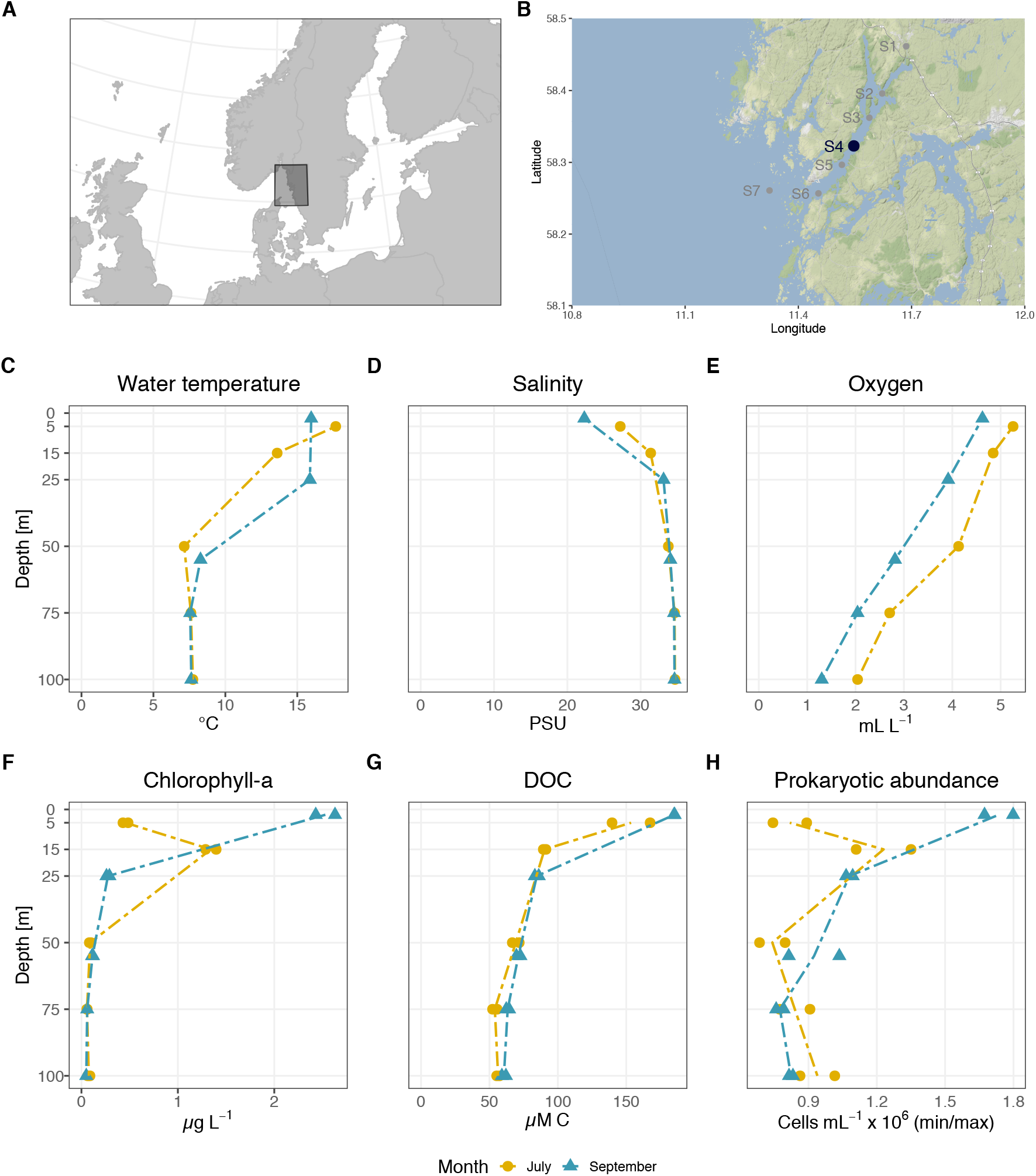
Overview of study site and water column characteristics. Location of the Gullmar Fjord on the Swedish west coast (**A**), and the sampling site at station 4 (Alsbäck) with a bottom depth of 120 m (**B**). Depth profiles of temperature (**C**), salinity (**D**), oxygen levels (**E**), chlorophyll *a* (**F**), dissolved organic carbon (DOC) (**G**), and prokaryotic abundance (**H**) in July and September.

### Nutrients, chlorophyll a concentration and prokaryote abundance

*In situ* nutrient concentrations were measured with a QuAATtro AutoAnalyzer and XY-3 Sampler (Seal Analytics) at the Sven Lovén Centre for Marine Infrastructure in Kristineberg, Sweden. Samples of 12 mL were poured in 13 mL polystyrene tubes (Sarstedt, 55.459) prior to the analysis of NO_3_^+^+NO_2_^+^ and, NH_4_^+^, PO_4_^3-^, and SiO_2_. For total nitrogen (TotN) and total phosphorus (TotP), 20 mL aliquots were analyzed with a QuAATtro AutoAnalyzer and XY-3 Sampler (Seal Analytics) with the protocol Q126 R0 Nitrate in seawater MT3B and Q125 R0 Phosphate in seawater MT3A. SiO_2_ was measured with the protocol NIOZ - Kisel Z06605_3 and NH_4_^+^ with Q033 R7 Ammonia MT3B.

Samples for determining DOC concentrations were filtered through 0.2 µm syringe filters (Acrodisc^®^ syringe filters, 32 mm, 514-4131, VWR), acidified with 448 μL of 1.2 M HCl to a pH of ~2 and analyzed with a high-temperature carbon analyzer (Shimadzu TOC-5000) at the intercalibrated facility at Umeå Marine Science Centre, Umeå, Sweden. DOC concentrations were calculated as non-purgeable organic carbon. Chl *a* concentrations were determined fluorometrically upon ethanol extraction as described in (24).

Samples for prokaryote abundance were fixed with 1% paraformaldehyde and 0.05% glutaraldehyde (final), stained with SYBR Green I nucleic acid stain (Invitrogen) (5 μM final concentration), spiked with 2 μL Flow Check High Intensity Green Alignment beads (Polysciences Inc.), and analyzed with a CyFlow Cube 8 flow cytometer (Sysmex Partec) according to (25) (Supplementary File 1).

### Metatranscriptomics analysis and sequence data accession

Water samples for metatranscriptomics were retrieved from every depth layer in biological duplicates. Approximately 3.5 L of water was filtered through Sterivex filter units (GP 0.22 μm, EMD Millipore) onboard within ~30 min after sampling, preserved in 2 mL RNAlater (Qiagen), immediately flash frozen in liquid nitrogen until samples were transferred to a −80°C freezer at the Sven Loven Center (Kristineberg, Sweden) and later stored at −80°C at Linnaeus University (Kalmar, Sweden) until further processing. Total RNA was extracted with RNeasy Mini Kit (Qiagen) according to (26) with some modifications as described in (27). In brief, total RNA was DNase treated using TURBO DNA-free Kit (ThermoFisher Scientific) and afterwards controlled for residual DNA by a 30-cycle PCR with 16S rDNA primers (27F and 1492R) including milli-Q as negative and DNA from *E. coli* as positive controls. Ribosomal RNA was depleted using RiboMinus Transcriptome Isolation Kit and RiboMinus Concentration Module (ThermoFisher Scientific). The rRNA-depleted fraction was linearly amplified using the MessageAmp II-Bacteria RNA Amplification Kit (ThermoFisher Scientific) before library construction (TruSeq) and sequencing at the National Genome Infrastructure, SciLifeLab Stockholm (Illumina HiSeq 2500 platform in rapid mode and with HiSeq SBS kit v4 chemistry to obtain 2 × 126 bp long paired-end reads) (Supplementary File 1).

Illumina adapter sequences were removed from quality controlled paired-end reads with Cutadapt (28) (v1.13) and reads quality trimmed with Sickle (https://github.com/najoshi/sickle) (v1.33) in paired end mode and sanger quality values. Ribosomal RNA sequences were removed by aligning them with ERNE (29) (v2.1.1) against an in-house database of stable RNA sequences from marine microbes. Next, rRNA filtered high quality reads were de-novo assembled with MEGAHIT (30) (v1.1.2) and default parameters. Subsequently, open reading frames (ORFs) were determined with Prodigal (31) (v2.6.3) and default parameters. Finally, the resulting peptides were used for a blastp search in the NCBI Refseq protein database with DIAMOND (32) (v0.9.24) and with an e-value threshold of 0.001. Taxonomic annotations were performed with MEGAN (33) (v6.12.8) with the additional longReads lcaAlgorithm for assemblies. Reads were mapped with bowtie2 (34) (v2.3.5.1) in paired end mode to the ORFs. Subsequently, Samtools (35) (v1.9) was used to quantify transcript counts per gene.

CAZyme ORFs were detected and classified with the run-dbcan program (36) (v2.0.11) with default parameters. PEPs were identified using HMMER3 (37) and Pfam profiles matching hits to PEP subunit sequences in the MEROPS database (https://www.ebi.ac.uk/merops). Transporter genes were detected and classified with HMMER3 using profiles from the Transporter Classification databases (TCDB). We noticed that the two PFAM domains PF00909 and PF00654 were assigned to incorrect TCDB families, therefore we manually reassigned the protein domains PF00909 to the “The Ammonium Channel Transporter (Amt) Family; TC 1.A.11” and PF00654 to the “The Chloride Carrier/Channel (ClC) Family; 2.A.49”. This incorrect assignment is noted in the TCDB database (Supplementary File 1).

The Thaumarchaeota-specific marker genes, RadA and AmoA peptides were detected by hidden Markov models run with HMMER3 and the profiles TIGR02236 (RadA) and PF12942 (AmoA), in the TIGRFAM (v15.0) and Protein Families (Pfam) (v31.0) databases. Hits were considered valid if the score was equal to, or higher than, the recommended “gathering score” for the model. In the case of 4-hydroxybutyryl-CoA dehydratase, an alignment was constructed using MUSCLE (38) with the peptide from the characterized gene in *Nitrosopumilus maritimus* SCM1 (locustag Nmar_0207), its predicted orthologs in other Thaumarchaeota (6) and all peptides that contained the two conserved domains according to their PFAMs in Nmar_0207, PF11794 (4-hydroxyphenylacetate 3-hydroxylase N terminal) and PF03241 (4-hydroxyphenylacetate 3-hydroxylase C terminal) in the set of peptides derived from the genomes in the MAR database using the “gathering score” as cutoff. A maximum likelihood phylogeny of the peptides was estimated using FastTree (39). Nmar_0207 clustered together away from predicted paralogs (peptides with a different function). To quantify Nmar_0207 ortholog expression, the peptides from the assembled metatranscriptome sequences were blasted using blastp with a relaxed e-value (0.0001) against the database of all MAR peptides together with Nmar_0207 predicted orthologs. Those whose closest hits were to Nmar_0207 orthologs were chosen to align them and place the new peptides on the phylogenetic tree. The packages PaPaRa (40), EPA-ng (41) and gappa (42) were used to confirm the position of the new peptides on the phylogenetic tree inside the group of 4-hydroxybutyryl-CoA dehydratase orthologs after visualization of the tree with iToL (43). The amino acid substitution model was predicted with IQ-TREE (44) (Supplementary File 1). All metatranscriptome data are available at the EMBL-EBI European Nucleotide Archive repository (https://www.ebi.ac.uk/ena), under the project accession PRJEB42919.

### Statistics and visualization

Principal component analysis (PCA) was performed on Hellinger transformed raw counts according to (45) on open reading frames (ORFs) with at least 5 cpm in more than 2 samples. Redundancy analysis (RDA) was performed on the same input data as described above. Environmental variables were standardized with the function *decostand* and selected based on pairwise Pearson correlation coefficients < 0.9 and variance inflation factors < 10. NH_4_^+^ concentrations below the detection limit of 0.2 μM were replaced with a small value (0.001) to enable the estimation of this variable in the tb-RDA. The suitability of an RDA was tested prior to analysis (gradient length of ~ 3.5). The model consisting of the variables Temperature, DOC, Chl *a*, NH_4_^+^, NO_3_^+^+NO_2_^+^, and PO_4_^+^ was significant (*p* < 0.001, *R^2^_adj_* = 57%). Monte Carlo permutation tests showed that i) both RDA axes were significant after Holm correction for multiple testing (*p_adj_* < 0.006) and ii) the variables DOC (*p_adj_* < 0.006), Chl *a* (*p_adj_* < 0.043), NH_4_^+^ (*p_adj_* < 0.04), NO_3_^+^+NO_2_^+^ (*p_adj_* < 0.043) were significant, explaining ~12% and ~9%, ~13, and 8% of the variation in prokaryotic community transcripts, respectively, as derived from the variance partitioning (Fig. S5). Grouping of samples was determined through hierarchical clustering based on scaled Hellinger transformed raw counts, Euclidean distances, and Ward D2 cluster criteria. The optimal number of clusters was graphically determined through Elbow and Silhouette methods. Observed richness (Chao1) and Evenness (Pielou’s J) were calculated with normalized counts that were scaled by ranked subsampling (SRS) according to (46). All statistical analyses and graphical visualization were conducted in RStudio (47) and predominantly with functions from the packages *tidyverse* (48) and *vegan* (49) (Supplementary File 1).

## RESULTS AND DISCUSSION

### Variability of biotic and abiotic parameters

Seawater samples were collected in duplicates from five depth layers (2-5, 15-25, 50-55, 75, and 100 m) in July and September 2017 at station Alsbäck (bottom depth ~120 m) in the Gullmar Fjord, Sweden (Fig. 1A, B). Analysis of CTD profiles showed a strong stratification of the water column both in July and September (Fig. 1C-E). The water temperature decreased rapidly from 13-17°C at the surface (2-25 m) to ~8°C from 50 m and below. In September, temperatures were similar at 16°C from 2 to 25 m and ~8°C from 50 m (Fig. 1C). The surface salinity in July was 28 PSU and decreased to 22 PSU in September, and at both times increased with depth to ~36 PSU at 100 m (Fig. D), which corresponds to North Sea values (50). Dissolved oxygen levels steadily decreased with depth from oxic (~4.5 to 5.2 mL L^−1^) at the surface to dysoxic (1 to 2 mL L^−1^) at the bottom (Fig. 1E). The vertical distribution of Chl *a* concentrations differed substantially between months. In July, Chl *a* at the surface reached ~0.5 μg L^−1^, increased to ~1.2 μg L^−1^ in the Chl *a* maximum at 15 m, and decreased to ~0.1 μg L^−1^ below 50 m depth (Fig. 1F). In September, concentrations were higher at the surface, reaching ~2.8 μg L^−1^ but rapidly decreased to below ~0.2 μg L^−1^ from 25 m downward (Fig. 1F). Interestingly, the variability between months seen for Chl *a*, was not reflected in the distribution of DOC concentrations, which remained at maximum levels (~170 μM C) at the surface and decreased ~3-fold to 100 m (Fig. 1G). Inorganic nutrients like NO_3_^+^+NO_2_^+^ and PO_4_^3-^increased from <1 μM to elevated levels at depth (up to 40 and 4.6 μM, respectively), although for example total nitrogen (~35 μM) remained relatively constant (Fig. S1E). The distribution of bacterial abundance resembled the vertical distribution of Chl *a*. Accordingly, bacterial abundance in July was generally low at ~0.9 × 10^6^ cell mL^−1^ except for a peak at ~1.2 × 10^6^ cells mL^−1^ in the Chl *a* maximum at 15 m depth. In September, bacterial abundance was highest in the surface at ~1.8 × 10^6^ cells mL^−1^ and steadily decreased with depth to ~0.9 × 10^6^ cells mL^−1^ at 100 m (Fig. 1H).

In terms of physicochemical water column characteristics and nutrient dynamics, the Gullmar Fjord compares to other fjord systems (19, 20). The Chl *a* concentrations measured here emphasize mesotrophic nature of the Gullmar Fjord (22). The ~2-fold difference between Chl *a* in July and September was likely due to elevated grazing pressure in July (personal observation), in line with previous observations (22).

### Overall patterns in prokaryotic community transcription by depth and month

Bacterial and archaeal transcription changed with depth and between samplings (Fig. 2). A principal component analysis (PCA) separated the widely spread surface samples (down to 25 m) from a tight cluster of deeper samples (50-100 m) (Fig. 2A). Detailed hierarchical cluster analysis showed four distinct clusters consisting of a July surface (5-15 m), a September surface (12 - 25m depth), and a separate cluster consisting of a mix of deep-water samples (50 - 100 m depth) (Fig. S2A). Further, redundancy analysis (RDA) showed that the influence of sampling date (July or September) on bacterial and archaeal transcription was strongest in the surface layer and decreased with depth (Fig. 2A, B). These differences were mainly explained by NH_4_^+^ (13% of variation) and partly by Chl *a* (9%), whereas depth variation was predominantly driven by DOC (12%), and NO_3_^−^+NO_2_^−^ (8%) (Fig. 2B and Fig. S3). This decrease in variability of transcription with depth was recently demonstrated for genes and taxa that showed diel oscillation in the surface but essentially diminished with depth through strong attenuation of light (51). Here, we show that this variability in transcription also applies to other gene systems, including CAZymes, PEPs, and transporters (TPs).

**Figure 2.**
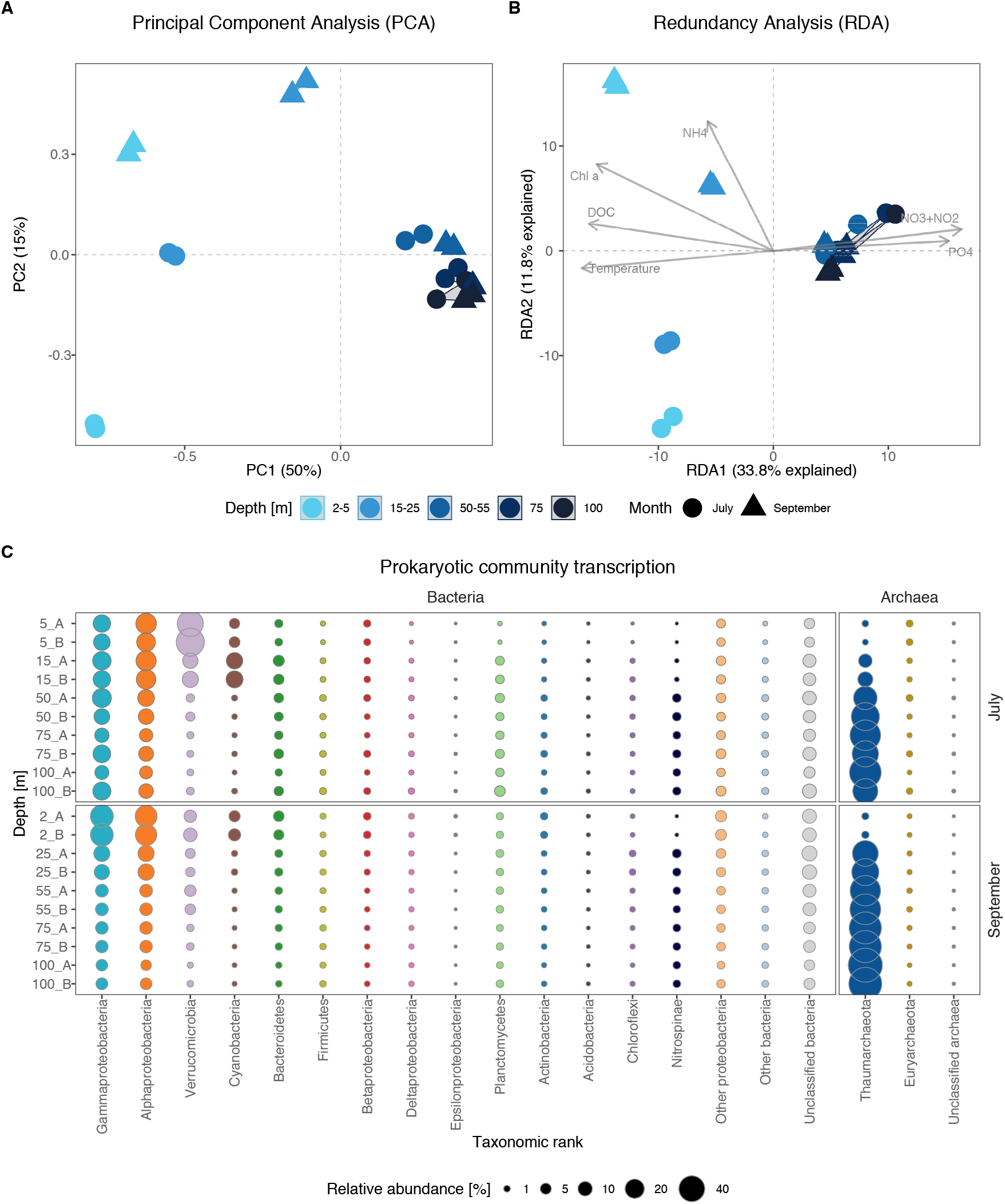
Variability in prokaryotic community transcription across depth and between sampling dates. Principal Component Analysis (PCA) based on 82842 quality filtered open reading frames (ORFs) (**A**). Redundancy Analysis (RDA) constrained by environmental variables (**B**). Taxonomic affiliation of the transcriptionally most abundant prokaryotic phyla (**C**). Note that Proteobacteria are shown at the class level.

Alpha- and Gammaproteobacteria accounted for high portions (around 25-30%) of the prokaryotic community transcription in the surface waters and maintained elevated transcription (~11%) throughout the water column (Fig. 2C). Also, Cyanobacteria (primarily *Synechococcus*) and Verrucomicrobia were highly active in the surface (reaching ~15% and 45% of community transcription, respectively), especially in September. These patterns are largely in agreement with marine metagenomic and metatranscriptomic surveys from the open ocean, which typically report a dominance of Picocyanobacteria (*Prochlorococcus*), Verrucomicrobia, Alphaproteobacteria (primarily the SAR11 clade), and Bacteroidetes in the upper water column (9, 18, 52). The relatively high activity of Verrucomicrobia in July is in line with a report from the Baltic Sea, where 16S rRNA gene analyses show that this taxon reaches higher relative abundances during summer coinciding with a dominance of *Cyanobacteria* (53). Metagenomic analysis of *Verrucomicrobia* shows that this taxon carries an extensive repertoire of CAZymes (e.g., alpha- and beta-galactosidases, xylanases, fucosidases, agarases, and endoglucanases), suggesting that they play a vital role in the degradation of phytoplankton-derived DOM and complex polysaccharides such as fucoidans (18, 54, 55).

Strikingly, Thaumarchaeota dominated transcription from 50 m downwards in July and from ~25 m in September, accounting for up to ~75% of total transcripts. Also, the functional contribution of Nitrospinae (~2.5%, <0.5% at 5 m) was noticeable in deeper water layers that were associated with lower oxygen concentrations (Fig. 2C and Fig. 1E). High proportions of Thaumarchaeota and Nitrospinae at depth have also been found in the northern Pacific and the Mediterranean Sea (9, 18, 52). Thaumarchaeota are key players in the oceanic N-cycle by oxidizing ammonia to nitrite (first step in nitrification) (6, 7) and generally increase in abundance with depth, accounting for up to 40% of total cells in the mesopelagic (56). Nitrospinae, in turn, are the most abundant and ubiquitously distributed nitrite-oxidizing bacteria (NOB) in the ocean (performing the second step in nitrification, oxidation of nitrite to nitrate) (8). In natural systems, Thaumarchaeota are typically ~10-fold more abundant than Nitrospinae (57). Thus, the transcriptional activity of Thaumarchaeota together with Nitrospinae below 50 m depth suggests an important contribution of these two groups to the N-cycle in this stratified fjord and potentially to the C-cycle since both taxa are lithoautotrophs.

### Variation in bulk prokaryotic transcription of DOM transformation genes (CAZymes, PEPs, and transporters)

Averaged over the entire water column, Bacteria accounted for 97% of CAZyme (~0.3% of total transcripts) and ~90% of PEP gene transcription (~3.4% of total transcripts). In contrast, Archaea devoted little transcriptional effort to CAZymes and PEPs, but instead accounted for ~55% of TP transcription (~10.4% of total transcripts). These patterns prompted us to determine the relationship between the three gene systems over depth (Fig. S8). The relative prokaryotic CAZyme transcription as compared to PEPs was up to five- to ten-fold higher in the surface waters than at depths from 50 m and below; note in particular the high proportion of CAZymes at 15 m in July (Fig. S4A). In addition, we found that the proportion of CAZyme and PEP transcription in relation to TPs decreased with depth (Fig. S4B and S4C), mainly because of the increasing dominance of Archaea. The higher proportion of CAZymes in the Chl *a* maximum layer agrees with enzymatic activity dynamics during phytoplankton blooms, particularly during bloom senescence, and in Chl *a* maximum layers in marine and limnic systems (10, 58). If the concentration of cleavage end-products determine prokaryotic ectoenzyme activities, the relative increase in PEP transcription with depth suggests that prokaryotic communities increase their efforts to acquire organic N and P because exported DOM typically has high C:N ratios (59). This has been shown for cell-specific enzymes such as leucine aminopeptidases and alkaline phosphatases, which typically increase in activity with depth (11).

These pronounced patterns in depth distributions, in turn, inspired further analysis of the allocation of transcriptional efforts to CAZymes, PEPs, and TPs among the dominant prokaryotic orders (Fig. S5). This highlighted the disproportionate contribution of Thaumarchaeota in nutrient uptake over polymer degradation at depth. Alphaproteobacteria such as Pelagibacterales showed a relatively high and stable proportion of TP transcripts compared to degrading enzymes with depth and across sampling dates. A somewhat different trend was noticed for Rhodobacterales that potentially engaged more in carbohydrate degradation in the upper water column but changed toward proteins with depth; this was more pronounced in September than in July (Fig. S5). Interestingly, a preference for carbohydrates relative to protein was more accentuated in Flavobacteriales in the upper water column but shifted toward PEPs with depth, whereas Cellvibrionales invested consistently more in CAZymes than PEPs throughout the water column in July but showed a similar trend to Flavobacteriales in September, emphasizing their significant role in the turnover of carbohydrates in surface waters. Alteromonadales, in turn, contributed more to the expression of TPs relative to CAZymes and seemingly favored peptides over carbohydrates in subsurface layers (Fig. S5). Synechococcales showed a disproportionate investment in TPs compared to degrading enzymes from 2 to 15 m depth. However, PEPs and TPs increased over CAZymes from 50 m downwards with a disproportionate transcription of PEPs over TPs.

These results suggest that the prokaryotic community attempted to acquire N and P with depth, as seen by the disproportional investment in PEPs and TPs over CAZymes. However, the relative proportions of these gene systems varied substantially between orders throughout the water column and were surprisingly consistent between sampling dates for some orders (e.g., Thaumarchaeota, Pelagibacterales, and Alteromonadales). In addition, our findings show that a few bacterial groups, such as Bacteroidetes, Cellvibrionales, Planctomycetes, and Verrucomicrobia showed relatively higher transcription of CAZymes compared to transporters - particularly in the surface, suggesting that marine microbes regulate their transcriptional efforts in degrading enzymes and transporters in a depth-dependent manner. The variation in the ratios implies a crucial role of different bacterial and archaeal taxa in the processing of HMW-DOM emphasizing their metabolic/transcriptional plasticity in transcribing CAZymes, PEPs, and TPs. Possible triggers of such variation could be changing elemental ratios of DOM and the desire to maintain a balanced stoichiometry. If confirmed in future analysis, this would have implications for the constraints of DOM cycling and microbial population dynamics.

### Divergence in CAZyme transcription

The richness of expressed CAZymes differed significantly between different depth layers (ANOVA; *F*_4,15_ = 14.87, *p* < 0.00004) (Fig. 3A), with ~4-fold higher values in the surface (2-5 m depth) compared to the deeper water layers (15-100 m depth) (Tukey; *p* < 0.03 - 0.0001). While both richness (Fig. 3A) and evenness (Fig. 3B) were fairly similar in July, the evenness in September at 25 m was lower compared to the deep.

**Figure 3.**
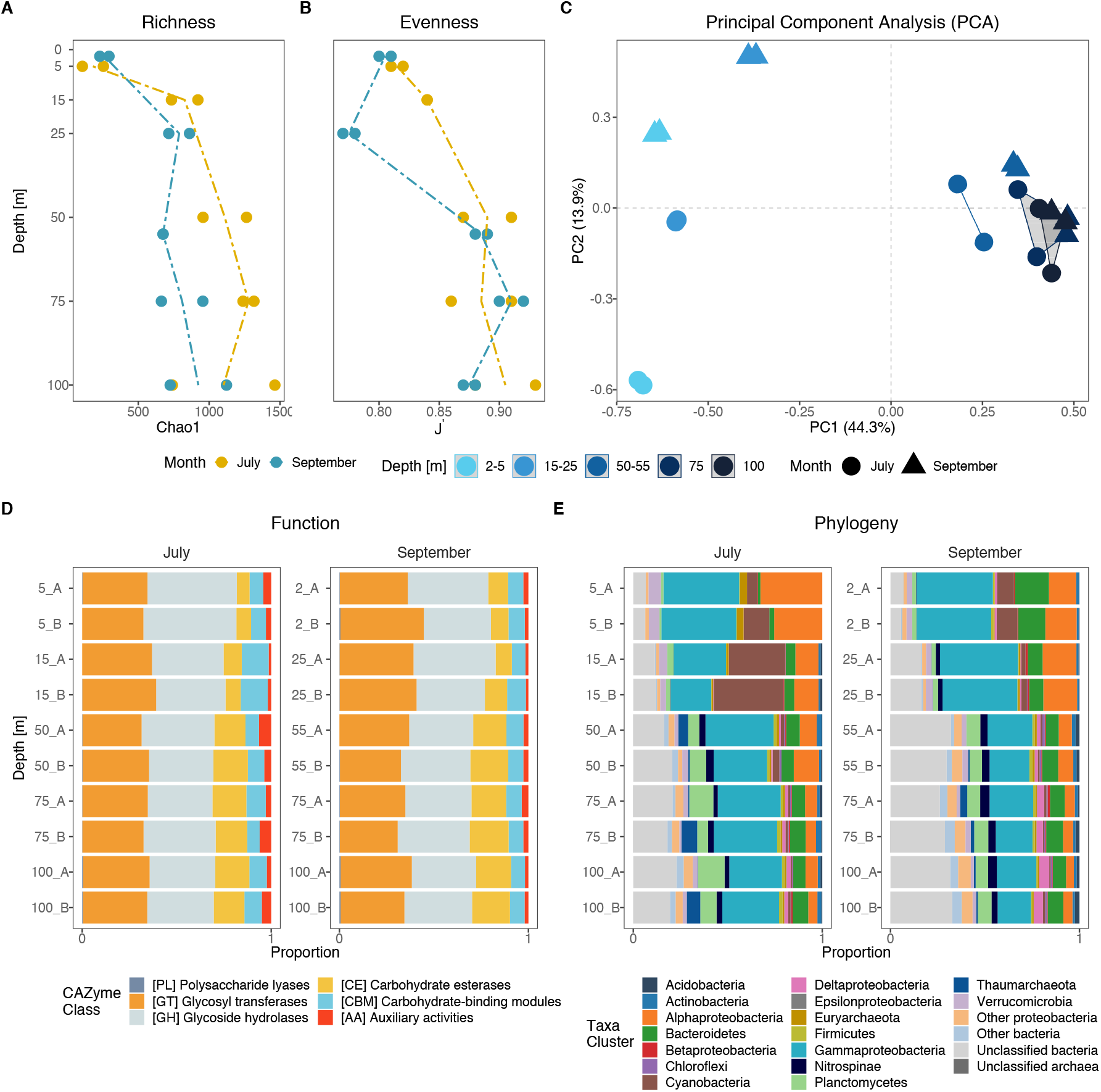
Taxonomic and functional profiles of carbohydrate-active enzyme (CAZyme) transcripts. The estimated richness (Chao1) (**A**) and evenness (J’) (**B**) depth profiles of CAZymes at the ORF level. Principal Component Analysis (PCA) of 1350 CAZyme affiliated open reading frames (ORFs) (**C**). Functional transcripts at the CAZyme class level (**D**), and taxonomic affiliation of the most active phyla transcribing CAZymes (**E**). Proteobacteria are grouped at the class level.

The microbial community expression of CAZyme genes differed significantly with both depth (PERMANOVA, *R^2^* = 0.61, *p_adj_* < 0.0003) and month (PERMANOVA, *R^2^* = 0.1, *p_adj_* < 0.001) (Fig. 3C). The CAZyme transcription in the surface layer (2-5 m) clustered with the samples from 15-25 m, and distantly from the deeper samples (50 - 100 m) (Fig. S2C). Differences in clustering of samples between July and September were observed for the surface samples (2-5 and 15-25 m), but only minimally for the samples below 50 m depth (Fig. 3C and Fig. S2C).

The relative proportions of CAZyme classes were relatively stable throughout the water column (Fig. 3D), with the most abundant classes being glycoside hydrolases (GHs: 37.9 ± 4.4%, n = 20) and glycosyltransferases (GTs: 35.6 ± 3.4%). This was surprising given that the taxonomic affiliation of these genes changed with depth (Fig. 3E). Nevertheless, Zhao et al. (2020) found a similar pattern among epi- to bathypelagic samples from the Pacific, Atlantic and Southern Ocean (16). This suggests that the CAZyme class level is too coarse to identify functional changes across depths, which is in stark contrast to the distribution of CAZyme families (see below; Fig. 3 and 4).

**Figure 4.**
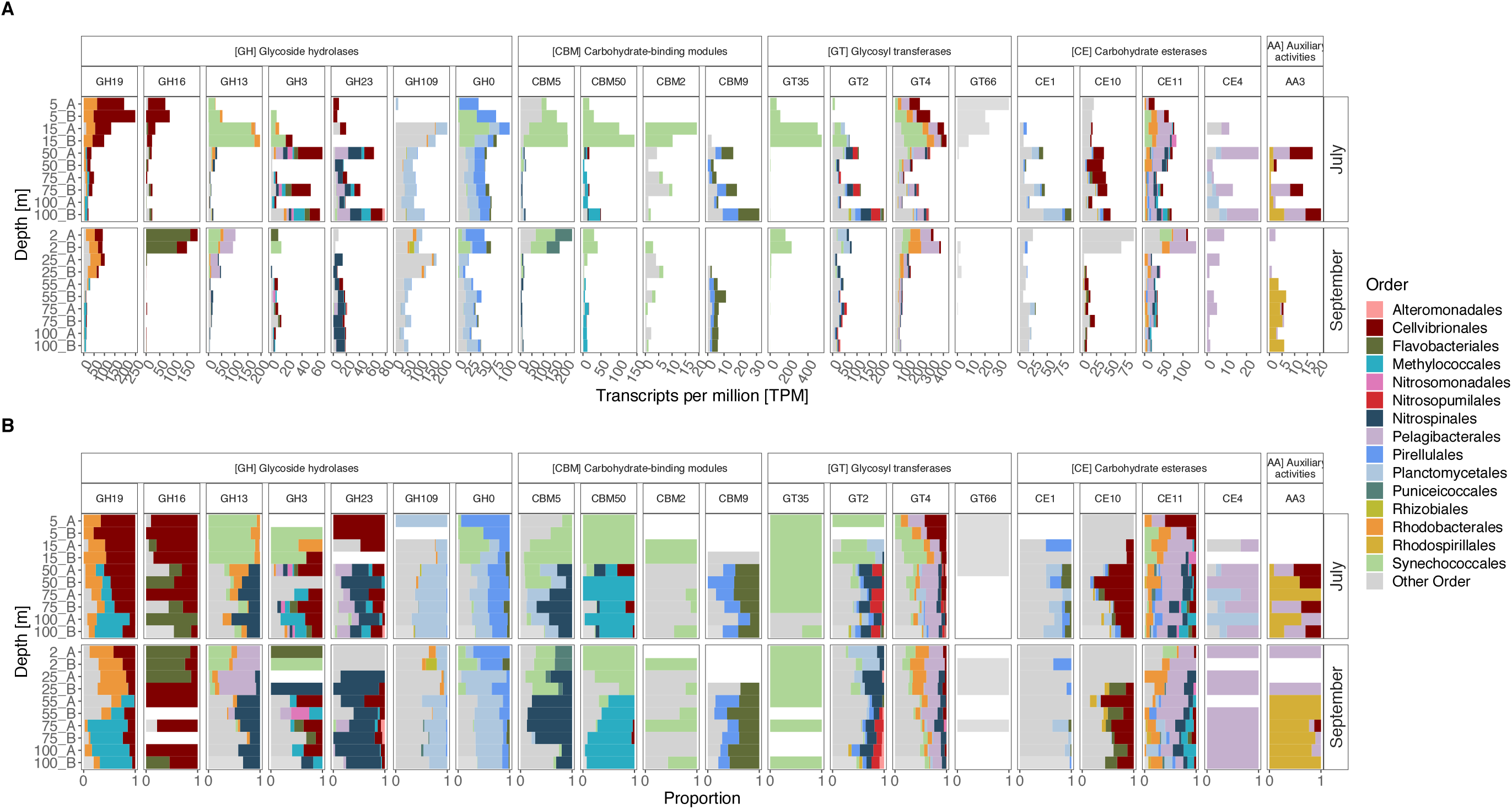
Transcription of the 20 most abundant carbohydrate-active enzymes (CAZymes) grouped into CAZyme families and taxonomic orders. Relative transcript abundance in transcripts per million (TPM) (**A**); note different scales on Y-axes. Proportions of CAZyme transcripts (**B**). Individual CAZyme families are further grouped into CAZyme classes (top facets).

Gammaproteobacteria dominated CAZyme expression both in July and September, accounting for up to ~40% of CAZymes in the upper water layers and 10-30% from 50 m and below (Fig. 3E). Alphaproteobacteria were also abundant in transcription in the surface layers (up to ~30%), with expression levels down to 5% of CAZymes at depth. Interestingly, Cyanobacteria dominated in July in the subsurface Chl *a* maximum layer where they accounted for ~34% of total CAZyme transcription. The significantly lower evenness observed in September (Fig. 3B) was likely the result of the Gammaproteobacteria being composed of a diverse set of taxa with a more uneven distribution of transcripts (Fig. 3B). Notably, from 50 m downward, Planctomycetes, Nitrospinae, and Deltaproteobacteria accounted for ~8%, 3.6%, and ~2.6% (both months) of total CAZyme transcription, respectively (Fig. 3E). Verrucomicrobia showed a relatively stable activity profile with depth, contributing 1-6% of CAZyme expression.

At the order level, pronounced differences in the most abundant CAZyme families between months and with depth (Fig. 4 and Fig. S6). For instance, Cellvibrionales and Rhodobacterales dominated transcription of glycoside hydrolase family 19 (GH19) in the surface layer, whereas Methylococcales dominated below 50 m depth. Actually, GH19 explained 18.5% of the variation in community transcription with depth and ~27% by month (Table S1). GH19 primarily consists of chitinases that hydrolyze chitin, the primary component of cell walls in fungi and exoskeletons of crustaceans (60). Incidentally, we noticed high abundance of copepods in microscopy samples from July that coincided with the low Chl *a*, in line with previous observations from this time of year (22). These findings suggesting a crucial role of chitin and chitodextrins as resources for prokaryotic plankton. Cellvibrionales and Flavobacteriales both transcribed GH16, with a larger proportion of Flavobacteriales in September (Fig. 4). The GH16 family comprises laminarinases that allow bacteria to decompose the algal storage glucan laminarin (58). Cellvibrionales and Flavobacteriales are typically abundant and active during phytoplankton blooms and likely fulfil different roles in the degradation of phytoplankton-derived organic matter (58).

Synechococcales dominated transcription of GH13 at 15 m in July, with some contribution also from Alphaproteobacteria (e.g., Rhodobacterales and Pelagibactrales) (Fig. 4). GH13 contains for example alpha-amylases that allow hydrolysis of the storage polymers glycogen and starch (61). In addition, Synechcoccales showed a high transcription of glycosyltransferases (GT35 and GT4) and carbohydrate-binding modules (CBM50 and CBM5). The glycosyltransferase GT35 is associated with the synthesis of starch and glycogen (62), whereas GT4 is involved in the synthesis of cellular structures and energy storage (i.e., sucrose, mannose, and trehalose synthase) upon photosynthesis (15, 62). The carbohydrate-binding module CBM50 and CBM5 are involved in chitin binding to facilitate the degradation of chitin or peptidoglycan (63). Interestingly, macroalgae and cyanobacteria express an extensive suite of CAZymes (e.g., cellulases, amylases, galactosidases), PEPs, and lipases (64). Given that Cyanobacteria, including Synechococcales, synthesize the polysaccharide glycogen as an internal carbon and energy storage compound (65), the relatively high expression of GH13 is likely associated with the utilization of internal glycogen sources rather than being a sign of extracellular degradation. Indeed, cyanobacteria genomes encode a relatively low number of genes for secretory enzymes (16). The strong transcriptional response in CAZyme transcription by Synechococcales highlights the importance of these genes in regulating glycogen metabolism across pronounced light and nutrient gradients by balancing organic carbon synthesis and degradation.

Two taxa stood out as having higher CAZyme transcription from 50 m and down: Planctomycetales and Nitrospinales (Fig. 3). Planctomycetales dominated transcription of α-N-acetylgalactosaminidase (GH109) and glycoside hydrolases (G0), which are not yet classified in the CAZy database (Fig. 4). GH109 is involved in the degradation of bacterial cell walls (66) and has been shown to be abundant in metagenome-assembled genomes (MAGs) throughout the water column in the Mediterranean Sea (18) (Fig. 4). Since the cell walls of Planctomycetales lack the polymer peptidoglycan (67), the transcription of GH109 suggests a role in the degradation of cell walls of other bacteria. Nitrospinales increased transcription of GH23, CBM5, and GT2 with depth (Fig. 4). These CAZymes are involved in the degradation of peptidoglycan or chitin and the synthesis of chitin or cellulose (62, 66).

The transcriptional CAZyme responses by a diverse microbial community confirm the crucial role of the storage polysaccharide laminarin in fueling the heterotrophic carbon demand in this stratified fjord. Moreover, we noted strong transcriptional responses associated with the structural polysaccharides chitin and peptidoglycan. Remarkably, our results highlight a depth-layer dependent functional partitioning in CAZyme transcription by well-known polymer degraders like Cellvibrionales and Flavobacteriales. Importantly, our analysis identified taxa like Nitrospinales, Nitrosomonadales, and Methylococcales that previously seem to have been overlooked in the context of polysaccharide degradation.

### Divergence in membrane transporter transcription

The estimated richness of transcribed transporters increased with depth in July but was relatively constant in September (Fig. 5A). The evenness of transcribed transporters, however, was lower below 20 m depth, especially in September (Fig. 5B). Moreover, transcription of prokaryotic membrane transporter genes differed significantly with depth (PERMANOVA: *R^2^* = 0.62, *p_adj_* < 0.0003) and sampling date (PERMANOVA: *R^2^* = 0.08, *p_adj_* < 0.002) (Fig. 5C). As for overall transcription, cluster analysis of transporter transcription grouped samples into four distinct depth clusters (Fig. S2D). The largest variability in transcription was associated with the upper water layers, which clustered away from the samples from 50 m and below.

**Figure 5.**
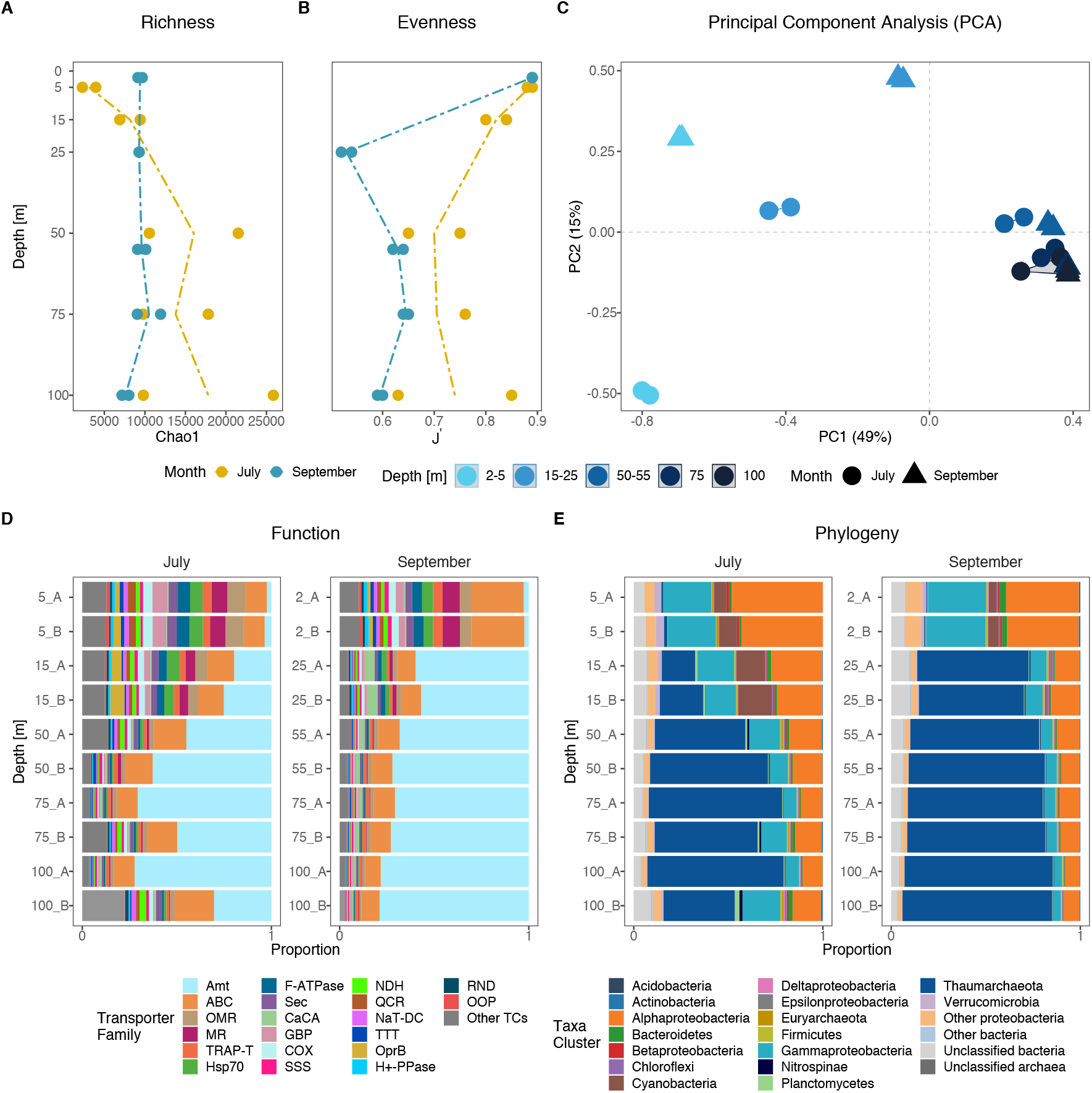
Taxonomic and functional transcription of membrane transporters (TPs). The estimated richness (Chao1) (**A**) and panel evenness (J’) (**B**) depth profiles of TPs at the ORF level. PCA of 25395 TP affiliated open reading frames (ORFs) (**C**). Functional transcripts of the most abundant TP families (**D**), and taxonomic affiliation of the most active phyla transcribing TPs (**E**); Proteobacteria are shown at the class level. Abbreviations of TC families: Amt - Ammonium Transporter Channel; ABC - ATP-binding Cassette; OMR - Outer Membrane Receptor; MR - Ion-translocating Microbial Rhodopsin; TRAP-T - Tripartite ATP-independent Periplasmic Transporter; Hsp70 - Cation Channel-forming Heat Shock Protein-70; F-ATPase - H^+^- or Na+-translocating F-type, V-type and A-type ATPase; Sec - General Secretory Pathway; CaCA - Ca^2+^:Cation Antiporter; GBP - General Bacterial Porin; COX - H^+^-translocating Cytochrome Oxidase; SSS - Solute:Sodium Symporter; NDH - H^+^ or Na^+^-translocating NADH Dehydrogenase; QCR - H^+^-translocating Quinol:Cytochrome c Reductase; NaT-DC - Na^+^-transporting Carboxylic Acid Decarboxylase; TTT - Tricarboxylate Transporter; OprB - Glucose-selective Porin; H^+^-PPase - H^+^, Na^+^-translocating Pyrophosphatase; RND - Resistance-Nodulation-Cell Division; OOP - OmpA-OmpF Porin.

The most abundant transporter families were ammonium transporters (Amt: 47 ± 28%, number of samples n = 20), ATP-binding Cassette transporters (ABC: 14 ± 6%, n = 20), and Outer Membrane Receptors (OMR: 4 ± 3%, n = 20) (Fig. 5D). PCA analysis of transporter transcription showed that these transporters contributed between 38% and 12% of variation in community gene expression with depth (PC1) and between 27% and 11% of variation by sampling date (PC2) (Table S1). Generally, transporter transcription in the surface was taxonomically similar to CAZymes. However, from 15 m depth and below, we noticed an exceptionally high contribution of Thaumarchaeota on both samplings, accounting for up to 79% of transporter transcription (Fig. 5E).

Our findings on membrane transporter transcription indicated that Pelagibacterales and Rhodobacterales were actively engaged in the turnover of low molecular weight compounds but depicted different nutritional preferences with noticeable changes between depths and sampling dates (Fig. 6 and Fig. S7). Both orders showed important contributions to e.g. Tripartite ATP-independent Periplasmic Transporter (TRAP-T), Tricarboxylate Transporter (TTT), and transporters for branched-chain amino acids (PF02653) and sugars (PF00532) (Fig. 6). However, pronounced differences in transcription in a variety of transporter families, including Solute:Sodium Symporter (SSS) and proteorhodopsin (microbial rhodopsin; MR, ~3% of transcription) were noted. For example, Rhodobacterales did not transcribe proteorhodopsins and showed a higher activity in TTT transcription in July compared to Pelagibacterales that contributed more in the surface in September (Fig. 6). Also, Pelagibacterales expressed high levels of glycine betaine transporters (PF04069), whereas Rhodobacterales transcribed more extracellular solute-binding proteins for amino acids, oligopeptides, and oligosaccharides (PF00497, PF00496, PF01547). The common osmolyte glycine betaine is readily degraded by marine bacteria including Pelagibacterales who require reduced sulfur compounds (e.g., dimethylsulfoniopropionate - DMSP or serine) for optimal growth (68, 69). Thus, they encode high affinity glycine betaine transporters which have a broad specificity for substrates with similar structures to glycine betaine such as DMSP, choline, and proline (70, 71). Similarly, for Alphaproteobacteria in the Atlantic Ocean, lowest relative proportions of transporter-related membrane proteins derive from Rhodobacterales in the mesopelagic (300 - 850 m), whereas Pelagibacterales show highest proportions in the euphotic zone down to the mesopelagic (72). Taken together, these differences in transcribed transporters together with differences in the depth distribution suggest pronounced functional partitioning of nutrients between Pelagibacterales and Rhodobacterales with depth.

**Figure 6.**
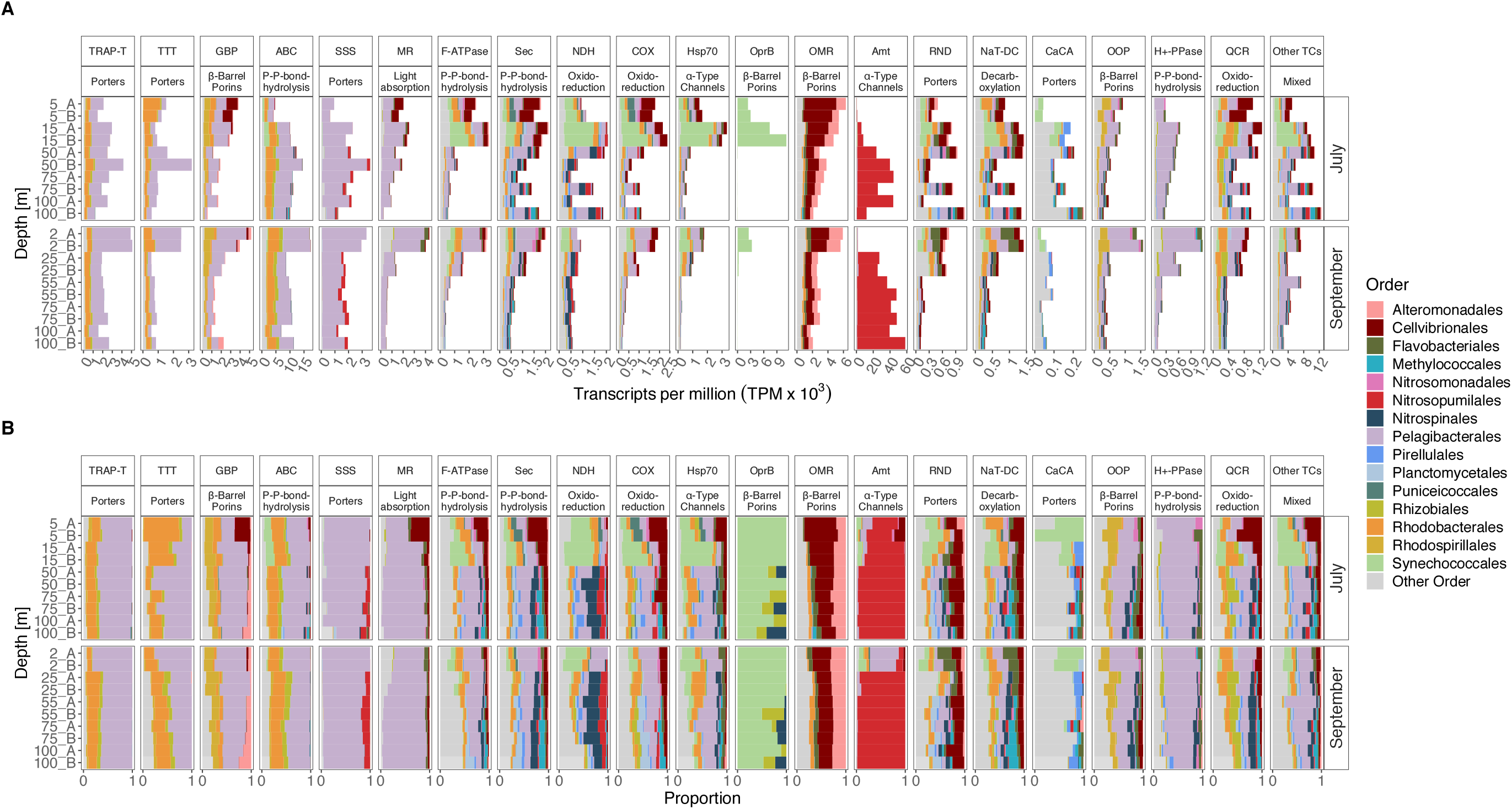
Transcription of the 20 most abundant transporter families (TPs) grouped into transporter subclasses and taxonomic orders. Relative transcript abundance in transcripts per million x 10^3^ (**A**); note different scales on Y-axes. Proportions of TP transcription of prokaryotic orders (**B**). Abbreviation of TC families are the same as in Fig. 5.

Pelagibacterales largely dominated the expression of proteorhodopsin (MR transporter family), along with some expression by Cellvibrionales and Flavobacteriales. As expected, proteorhodopsin expression was highest in the surface (especially in September), yet substantial expression by Pelagibacterales was detected also down to 100 m (i.e., well below the deep chlorophyll maximum at 25 m; Fig. 6). Proteorhodopsins (PR) are light-activated proton pumps that generate a proton motive force that in turn can be utilized by ATPases to generate energy (5). These retinal-binding rhodopsin proteins are widespread in bacteria (73, 74 and references therein) including Pelagibacterales (SAR11 clade) (69 and references therein). The precise metabolic processes that are stimulated by PR photoheterotrophy appear to differ between prokaryotes with different life strategies (73, 74). In Pelagibacterales, PR is constitutively expressed and under starvation conditions in the light, the bacteria can replace portions of the ATP production from respiration of DOC with ATP production from PR light harvesting (75). In this manner, PR conveys a competitive advantage under oligotrophic and mesotrophic conditions.

Synechococcales showed the highest transcript abundance of transporters in the subsurface Chl *a* maximum layer in July and to a lesser extent in September. Photoautotrophs which are distributed between the surface and the deep Chl *a* maximum experience sharp gradients in light and nutrients and therefore face the challenge to optimize growth by adapting their metabolic machinery (76). The transcriptional increase of ion and electron transporters (e.g., F-ATPase, Sec, NDH, COX, and Hsp70) in the Chl *a* maximum, together with the active usage of storage polysaccharides, indicates an active struggle to maintain a beneficial metabolic balance for growth, given their involvement in respiration and stress responses (77). Interestingly, Synechococcales dominated transcription of the carbohydrate-selective porin (OprB) family in July (in the Chl *a* maximum) (Fig. 6). To our knowledge, such high transcription of *oprB* by this taxon across a vertical depth profile has not been reported before. In *Pseudomonas putida,* these porins are part of a high affinity uptake system for a variety of carbohydrates, including glucose (78). Unicellular picocyanobacteria (*Prochlorococcus* and *Synechococcus*) also encode genes with an OprB domain (79–81) and uptake of glucosyl-glycerol, sucrose, and trehalose under temperature and salinity stress has been demonstrated (82 and references therein). Thus, picocyanobacteria may acquire a variety of low molecular weight carbohydrates through these porins either for osmotic adjustments (82) or in response to P starvation through uptake of sugar-phosphates (81). These results provide additional support for the importance of a certain capacity for mixotrophy in picocyanobacteria (83, 84).

An important transporter family dominated by Cellvibrionales and Alteromonadales (Gammaproteobacteria) was the OMR family, which reached the highest relative transcription in the surface layer and decreased with depth (Fig. 6). Although the relative transcription of OMR by Cellvibrionales was more than 3-fold higher than Alteromonadales in the July surface layer, the two taxa contributed equally in September. Transporters in the OMR family, which include TonB receptors, have broad substrate specificities and are involved in the uptake of iron-siderophore complexes, vitamin B12, nickel complexes, colicins, and carbohydrates (85). In the North Pacific Subtropical Gyre, Alteromonadales increased their transcriptional activity with depth (9) and showed the highest transporter protein abundance between 300-800 m depth in the Atlantic Ocean (9, 52, 72). Knowledge of the ecophysiology and ecology of Cellvibrionales is still scarce, but they appear to reach highest relative abundances in surface waters and coastal areas (86, 87). Here we showed that both bacterial groups dominated OMR transcription throughout the water column, suggesting a vital role of the two gammaproteobacterial orders in nutrient cycling (potentially related to carbohydrates).

Strikingly, below 15-25 m depth, there was a dramatic increase in ammonium transporter (Amt) family transcripts by Thaumarchaeota, accounting for up to ~79% of transporter transcripts (~22% of total transcripts) at 100 m depth (Fig. 2 and Fig. 6). Other transporter families transcribed by Thaumarchaeota were sodium-solute (SSS) and H^+^ or Na^+^-translocating Carboxylic Acid Decarboxylase (NDH). Elevated transcription of the *amt* gene by Thaumarchaeota has been reported from the North Pacific Subtropical Gyre from 25 to 500 m and a high abundance of Amt transporter proteins has been reported from the Atlantic Ocean (52, 72). Thaumarchaeota are abundant and widespread in marine waters and sediments (56, 88, 89), and account for major proportions of ammonia oxidation in mesopelagic waters of the open ocean through the key enzyme ammonia monooxygenase (encoded by *amoABC*) (7). Thaumarchaeota genomes encode high-affinity ammonium transporters that enable them to outcompete ammonia-oxidizing bacteria (AOB) under low ambient ammonium concentrations (nM range) (6, 90). The variety of ABC type transporters (e.g., for amino acids, oligopeptides, phosphonates), urea transporters, and Solute:Sodium Symporter (SSS) family transporters (72, 91, 92) enables utilization of other nitrogen sources than ammonia (93) and supports previous findings that some Thaumarchaeota are capable of using inorganic and organic matter to supplement their metabolism (72, 91, 94).

Collectively, these findings suggest that depth strongly shapes the transcription of transporters, similarly to CAZymes. The most pronounced shift in transcription occurred between the surface layer and 50 m depth largely consisting of a pronounced increase in thaumarchaeal ammonium transporters. These diverse responses in different membrane transporter families, that were associated with different taxa, highlight the important contribution of a few functional groups (e.g., Pelagibacterales, Rhodobacterales, and Thaumarchaeota) to the turnover of nutrients (e.g., amino acids, carbohydrates, glycine betaine, DMSP, and ammonium) across depth gradients.

### Thaumarchaeal ammonia oxidation and carbon fixation

To follow up on the exceptionally high transcriptional activity of Thaumarchaeota below 25 m, we analyzed marker genes for thaumarchaeal energy and carbon metabolism (Fig. S9). This revealed elevated transcription both of the gene for the large subunit of archaeal ammonia monooxygenase (*amoA*) mediating the first step of nitrification and the gene for archaeal inorganic carbon fixation (*hcd*; 4-hydroxybutyryl-CoA dehydratase) in the hydroxypropionate/hydroxybutyrate (HP/HB) cycle (6). Relating this expression to the expression of the single copy gene *radA* (95) showed that the relative investment in ammonia oxidation was high (i.e., *amoA:radA* ratios 3-4-fold) compared to that of inorganic carbon fixation (4-hydroxybutyryl-CoA dehydratase gene:*radA* ratios ~ −3fold), especially in the upper surface layer (Fig. S8).

Although archaeal ammonia oxidation is coupled with inorganic carbon fixation (96), the low energy yield from ammonia oxidation (ΔG = −307.35 kJ mol^−1^ NH_3_) (97) suggests that a large quantity of ammonium needs to be oxidized per amount of CO_2_ (98). Accordingly, in soil Thaumarchaeota, the expression of *amoA* is several-fold higher than the *hcd* (99) and in *Nitrosopumilus adriaticus*, nitrification resulted in a C yield per N of ~0.1 (100). Our gene expression data extend these findings on cultivated isolates to the natural marine environment and suggest that different Thaumarchaeota may have a distinct influence on the oceanic nitrogen compared to carbon cycles.

## Conclusion

The overall depth distribution of prokaryotic transcription in the Gullmar Fjord agreed with findings in the Mediterranean Sea, North Pacific, and other fjords (18, 21, 52). Our findings provide novel insights into the stratification of transcriptional activity of genes involved in transformation of DOM and nutrient uptake in natural microbial communities, which heretofore primarily has been surveyed through metagenomics (16–18). The pronounced transcriptional differences in CAZymes among distinct taxa of bacteria and archaea between samplings and over depth were notable in several respects. Given the succession often observed among phytoplankton and in relation to zooplankton grazers, we infer that ecosystem level approaches are urgently needed to disentangle the scale of carbon fluxes associated with bacterial utilization of phytoplankton storage polysaccharides as compared to zooplankton structural polysaccharides. Moreover, building on previous studies into the genetic adaptations in nutrient scavenging with depth (72 and references therein), we suggest that expression analyses at the gene and protein levels are necessary to inform on the mechanisms regulating the spatial partitioning of resources between bacterial and archaeal taxa. Characterizing and quantifying the constraints on nutrient and carbon cycling in particular depth-layers, inhabited by specific sets of prokaryotic taxa, will be important for understanding how future changes in plankton dynamics will influence the efficiency of the biological carbon pump.

## ACKNOWLEDGEMENTS

We acknowledge the Sven Lovén Centre for Marine Sciences for their hospitality during our stay, in particular Peter Tiselius for his kind support with equipment, and Hans Olsson, Lars Ljungqvist, Bengt Lundve, and Pia Engström for their assistance throughout our stays regarding laboratory equipment and nutrient analyses. We also thank Ursula Schwarz and Carl Kristenssen for operating the R/V *Oscar von Sydow* and their skilful assistance during the field sampling campaigns. A special thank goes to Sabina Arnautovic and Camilla Karlsson for their dedicated work in the laboratory. This research was supported by a grant from the University of Gothenburg and the Royal Swedish Academy of Sciences (KVA) to BP, CB, and OCMG Support was also given through a grant from the Swedish Research Council VR to JP. CB was additionally supported by HIFMB, a collaboration between the Alfred-Wegener-Institute, Helmholtz-Center for Polar and Marine Research, and the Carl-von-Ossietzky University Oldenburg, initially funded by the Ministry for Science and Culture of Lower Saxony and the Volkswagen Foundation through the “Niedersächsisches Vorab” grant program (grant number ZN3285). JMG research was supported by the Spanish Ministry of Science and Innovation (project PID2019-110011RB-C32). The authors acknowledge support from the Science for Life Laboratory, the National Genomics Infrastructure, NGI, and Uppmax (compute project SNIC 2017/7-419 and storage project SNIC 2020/16-76), Sweden, for providing assistance in massive parallel sequencing and computational infrastructure.

## AUTHOR CONTRIBUTIONS

BP, CB, OCGM, and JP designed the study. BP, CB, OCMG, and CPM conducted the field work, retrieved samples, and processed samples in the laboratory. BP, DL, and JMG processed metatranscriptomic data. BP and JP wrote the manuscript with contribution from all authors.

